# Identification of high seed oil yield and high oleic acid content in brazilian germplasm of winter squash (*Cucurbita moschata* D.)

**DOI:** 10.1101/2021.07.23.453548

**Authors:** Ronaldo Silva Gomes, Ronaldo Machado Júnior, Cleverson Freitas de Almeida, Rebeca Lourenço de Oliveira, Rafael Ravaneli Chagas, Ednângelo Duarte Pereira, Fabio Teixeira Delazari, Derly José Henriques da Silva

**Author notes:** **Corresponding author**: PH Rolfs Avenue, Genetic Resources Management Laboratory, Zip Code 36570-900, Viçosa, MG, Brazil.

## Abstract

*Cucurbita moschata* D. seed oil contains approximately 75% unsaturated fatty acids, with high levels of monounsaturated fatty acids and antioxidant compounds such as vitamin E and carotenoid, constituting a promising food in nutritional terms. Associated to this, the Brazilian germplasm of *C. moschata* exhibits remarkable variability, representing an important source for the genetic breeding of this vegetable and other cucurbits. In this context, the present study evaluated the productivity and profile of the seed oil of 91 *C. moschata* accessions from different regions of Brazil and maintained in the Vegetable Germplasm Bank of the Federal University of Viçosa (BGH-UFV). A field experiment was conducted between January and July 2016. The tested *C. moschata* accessions showed high genetic variability in terms of characteristics related to seed oil productivity (SOP), such as the mass of seeds per fruit and productivity of seeds, providing predicted selection gains of 29.39 g and 0.26 t ha^-1^, respectively. Based on the phenotypic and genotypic correlations, greater SOP can be achieved while maintaining high oleic acid content and low linoleic acid content, providing oil of better nutritional and chemical quality. In variability analysis, the accessions were clustered into five groups, which presented different averages for SOP and fatty acid content of seed oil; approach that will guide the use of appropriate germplasm in programs aimed at genetic breeding for SOP and seed oil profile. *Per se* analysis identified BGH-4610, BGH-5485A, BGH-6590, BGH-5556A, BGH-5472A, and BGH-5544A as the most promising accessions in terms of SOP, with average (μ+g) of approximately 0.20 t ha^-1^. The most promising accessions for higher oleic acid content of seed oil were BGH-5456A, BGH-3333A, BGH-5361A, BGH-5472A, BGH-5544A, BGH-5453A, and BGH-1749, with average (μ+g) of approximately 30%, and almost all of these accessions were also the most promising in terms of lower linoleic acid content of seed oil, with average (μ+g) of approximately 45%. Overall, part of the *C. moschata* accessions evaluated in the present study can serve as a promising resource in genetic breeding programs for SOP and fatty acid profile, aiming at the production of oil with better nutritional and physicochemical quality.

## 1. INTRODUCTION

Winter squash (*Cucurbita moschata* D.) is one of the cucurbit vegetables of great socioeconomic importance owing to the high nutritional value of its fruits and seeds. Cultivated mainly for fruit production, *C. moschata* has been strategically used in biofortification programs for vitamin A because of its high content of carotenoids in fruits such as *β*- and *α*- carotene—the major precursors of vitamin A (Carvalho et al., 2012; Saltzman et al., 2013). In addition, winter squash fruits are an excellent source of minerals such as K, Ca, P, Mg, and Cu (Nagar et al., 2018; Priori et al., 2018). *C. moschata* is cultivated across a wide geographical range worldwide, and together with other cucurbits such as *C. pepo* and *C. maxima*, the area under the cultivation and worldwide production of *C. moschata* was estimated to be nearly 2 million hectares and 27.6 million tons, respectively, in 2018 (FAO, 2020), highlighting the socioeconomic importance of this vegetable.

Furthermore, *C. moschata* seed oil can serve as an excellent product due to its nutritional and physicochemical properties, associated with the high seed production potential of this vegetable. Lipids account for up to 49% of *C. moschata* seed components (Jarret et al., 2013; Patel, 2013), and studies on the germplasm of this cucurbit have already identified accessions that can produce up to 0.58 t·ha^-1^ of seeds (Gomes et al., 2020). *C. moschata* seed oil contains approximately 75% unsaturated fatty acids, with high content of monounsaturated fatty acids (MUFAs), such as oleic acid (Jarret et al., 2013; Sobreira, 2013; Veronezi and Jorge 2015). Thus, it is an excellent substitute for vegetable lipid sources that contain high levels of saturated fatty acids, which are harmful to human health. Associated with this, some studies have reported that *C. moschata* seeds and seed oil contain high levels of antioxidant compounds, such as vitamin E and carotenoids, components beneficial to human health (Veronezi and Jorge 2012; Dash et al., 2017), which also protect the seed oil from oxidative processes that may lead to rancidity. In this line, *C. moschata* seed oil may serve as a health food in the cultivation regions of this vegetable, particularly in less developed regions, and in the family farming context (Gomes et al., 2020).

Studies on the Brazilian germplasm of *C. moschata* have emphasized the evaluation of agromorphological characteristics reporting remarkable variability in these traits (De Lima et al., 2016; Ferreira et al., 2016; Oliveira et al., 2016; Gomes et al., 2020). As an allogamous species, variability in the *C. moschata* germplasm is associated with the occurrence of natural hybridization across different populations. Already present in the diet of native Latin American people (Dillehay et al., 2007; Piperno et al., 2003), and with a widespread cultivation in the American continent, the variability of *C. moschata* may also be related to anthropogenic actions, such as frequent exchange of seeds among family farmers (Gomes et al., 2020). Additionally, the variability of the Brazilian germplasm of *C. moschata* reflects its adaptation to a broad ecological range, constituting different edaphoclimatic conditions; thus, these accessions represent an important source for the genetic breeding of this vegetable and other cucurbits.

The Vegetable Germplasm Bank of the Federal University of Viçosa (BGH-UFV) maintains approximately 350 accessions of *C. moschata*, mostly landraces, with a collection period of over five decades, from different geographic regions of Brazil (Silva et al., 2001). The *C. moschata* collection maintained in the BGH-UFV constitutes a substantial sample of the Brazilian germplasm, being one of the largest collections of this species in the country (Fonseca et al., 2015). A preliminary assessment of the seed oil fatty acid profile of a small part of the *C. moschata* germplasm maintained in the BGH-UFV revealed high variability among 54 accessions in terms of oleic acid content (Sobreira, 2013). In that study, BGH-7765, with oil oleic acid content of 28.39%, was identified as a promising accession for use as parent germplasm in breeding programs aimed at improving *C. moschata* seed oil.

To this end, the objectives of the present study were (a) to evaluate the seed oil productivity (SOP) and oil fatty acid profiles of 91 *C. moschata* accessions from different regions of Brazil maintained in the BGH-UFV; (b) to analyze the correlations between these characteristics; and (c) to examine the variability of this germplasm for identifying accessions with high SOP and high oleic fatty acid content but low linoleic acid content in seed oil.

## 2. MATERIAL AND METHODS

### 2.1 Germplasm origin

The present study evaluated 91 *C. moschata* accessions maintained in the BGH-UFV. These accessions, mostly landraces, have been collected by the BGH-UFV over a period of more than five decades (Silva et al., 2001) from different geographical regions of Brazil (Figure 1). The accessions were evaluated with four genotypes used as controls, the cultivars ‘Jacarezinho’ and ‘Maranhão’ and the hybrids Jabras and Tetsukãbuto, which are widely cultivated and commercialized in Brazil.

**Figure 1.**
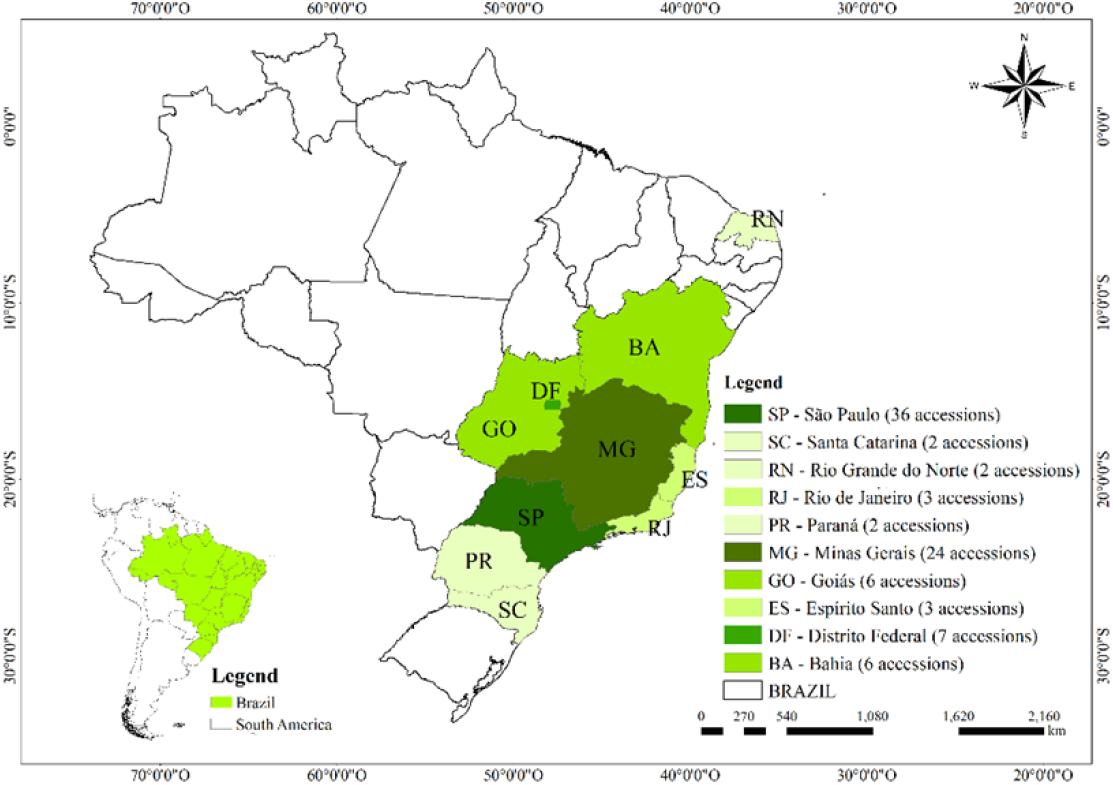
Brazilian map displaying the regions of origin of the *Cucurbita moschata* accessions tested in the present study.

### 2.2 Location and conduct of the experiment

A field experiment was conducted between January and July 2016 at “Horta Velha”— an experimental unit of the Department of Agronomy of UFV (20°45′24″S, 42°50′45″W; altitude, 648.74 m).

The genotype seedlings were cultivated in expanded polystyrene trays with 72 cells containing a commercial substrate. Subsequently, the seedlings were transplanted in the experimental area following to the augmented block design proposed by Federer (1956), with five replicates for each control. The plants were distributed with 3 × 3 m spacing between plants and rows, resulting in a stand of 1,111 plants ha^-1^. Each experimental plot contained five plants, and all evaluations of fruits, seeds, and seed oil were performed on three fruits each from three central plants in a plot.

### 2.3 Assessment of seed oil productivity

The genotypes were evaluated for the number of fruits per plant (NFP), total mass of seeds per fruit (MSF), productivity of seeds (PS), and total seed oil content (SOC). The PS, SOC, and SOP estimates were obtained using the following equations (equations 1, 2, and 3, respectively):

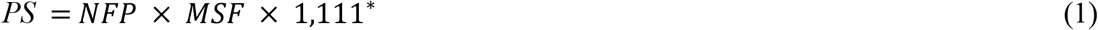

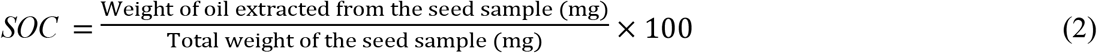

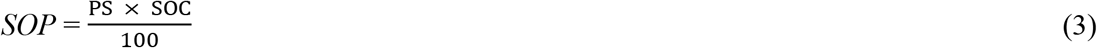

where PS is the productivity of seeds (t ha^-1^); NFP is the number of fruits per plant; MSF is the mass of seeds per fruit (g); ^*^1,111 is the number of plants per hectare; SOC is the oil content expressed as the percentage of the seed mass on a dry basis (%); and SOP is the seed oil productivity (t ha^-1^).

Initially, the seeds were dried in a forced air circulation oven for 72 h at 30°C. Next, 20 g seeds from each genotype were ground in a Willey knife mill with a 1 mm sieve. SOC was determined in an extractor (ANKOM XT15) following extraction with petroleum ether using a standard AOAC method (Thiex et al., 2003). Before loading in the extractor, the ground seed samples were dried in an unventilated oven at 105°C for 2 h. Then, approximately 2 g samples were transferred to filter envelopes (XT4; ANKOM technology), which were sealed and placed in the extractor. Oil was extracted from the samples with ether circulation for 30 min at 90°C, and the percentage of oil was calculated as the difference between the sample weight before and after extraction. SOC was expressed in grams per 100 grams of seeds on a dry basis.

### 2.4 Analysis of the fatty acid profile of seed oil

Seed oil extracted by cold pressing was subjected to gas chromatography. Oil was extracted from approximately 50 g of seeds using a hydraulic press aid according to the methodology described by Gomes et al. (2020).

The composition of the methyl esters of oil fatty acids was determined according to the methodology described by Bubeck et al. (1989), with some modifications. The Shimadzu GC-17A gas chromatograph equipped with an automatic insertion platform, a flame ionization detector, and a Carbowax capillary column (30 m × 0.25 nm) was used. Chromatography was performed at an injector temperature of 230°C and detector temperature of 250°C. The column operation was programmed to start at 200°C, with an increase of 3°C·min^-1^ until reaching the final temperature of 225°C. Nitrogen was used as the carrier gas at a flow rate of 1.3 L·min^-1^. The content of each methyl ester of fatty acids was expressed as a percentage of relative peak area.

### 2.5 Obtaining of best non-biased linear predictions (BLUPs), best non-biased linear estimates (BLUEs), variance components, and genetic-statistical parameters

Data were analyzed using a mixed model based on the predictions of restricted maximum likelihood (RML) and BLUPs. The “lme4” package in R version 3.6.1 was used (Bates et al., 2015). The variance components were obtained based on RML, and the genotypic values were obtained based on BLUPs and BLUEs using the following model:

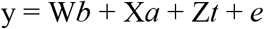

where y represents the vector comprising the phenotypic values of variables; *b* represents the vector comprising the effect of blocks (random effect); *a* represents the vector comprising the effect of accessions (random effect); *t* represents the vector comprising the control effect (fixed effect); and *e* represents the error vector. W, X, and Z represent the incidence matrices of parameters *b*, *a*, and *t*, respectively, with the data vector y. All statistical analyses were performed based on the genotypic values.

The genetic parameters were obtained based on the following estimators:

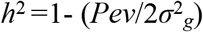

where *Pev* represents the prediction of error variance (Cullis et al., 2006)

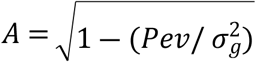

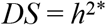

where *DS* represents the selection differential, estimated from an average of 15% of the most promising accessions for each characteristic.

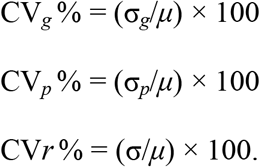

### 2.6 Analysis of correlations between characteristics

The correlations between characteristics were analyzed based on the following model:

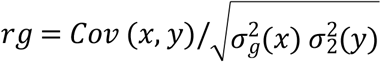

where *Cov* (*x, y*) represents the genetic covariance between two variables, X and Y, and σ^2^_*g*_ (*x*) and e σ^2^_*g*_ (*y*) represent the genetic variances corresponding to the variables X and Y, respectively.

The significance of correlations was analyzed in GENES (Cruz, 2013) using the Mantel test (Z statistic) at 1% and 5% probability.

### 2.7 Analysis of the germplasm variability

The matrix of distances between the genotypes was obtained based on the genotypic values of characteristics related to SOP and seed oil profile. The distances between genotypes were estimated as the average Euclidean distance with data standardization. Based on this, variability was assessed using Tocher’s clustering in GENES (Cruz, 2013).

Principal component analysis was used to identify the contribution of the characteristics to genotype variability using GENES (Cruz, 2013).

### 2.8 Identification of promising accessions

Promising accession clusters were identified, and *per se* analysis of the most promising accessions for each characteristic was performed. *Per se* identification was performed based on the ranking of the respective genotypic values and the environmental interaction-free genotypic value (μ+g), considering 15% of the most promising accessions.

## 3. RESULTS

### 3.1 Variance components and genetic-statistical parameters of characteristics related to SOP and fatty acid profiles

MSF was the characteristic associated with SOP with the highest genotypic variance (Table 1). Regarding oil fatty acid profile, oleic acid content exhibited the highest genotypic variance. Linoleic acid and polyunsaturated fatty acid (PUFA) content exhibited genotypic variances of 10.66% and 10.65%, respectively. All these variances were significant (*p* <0.01), as shown in Table 1.

**Table 1.**
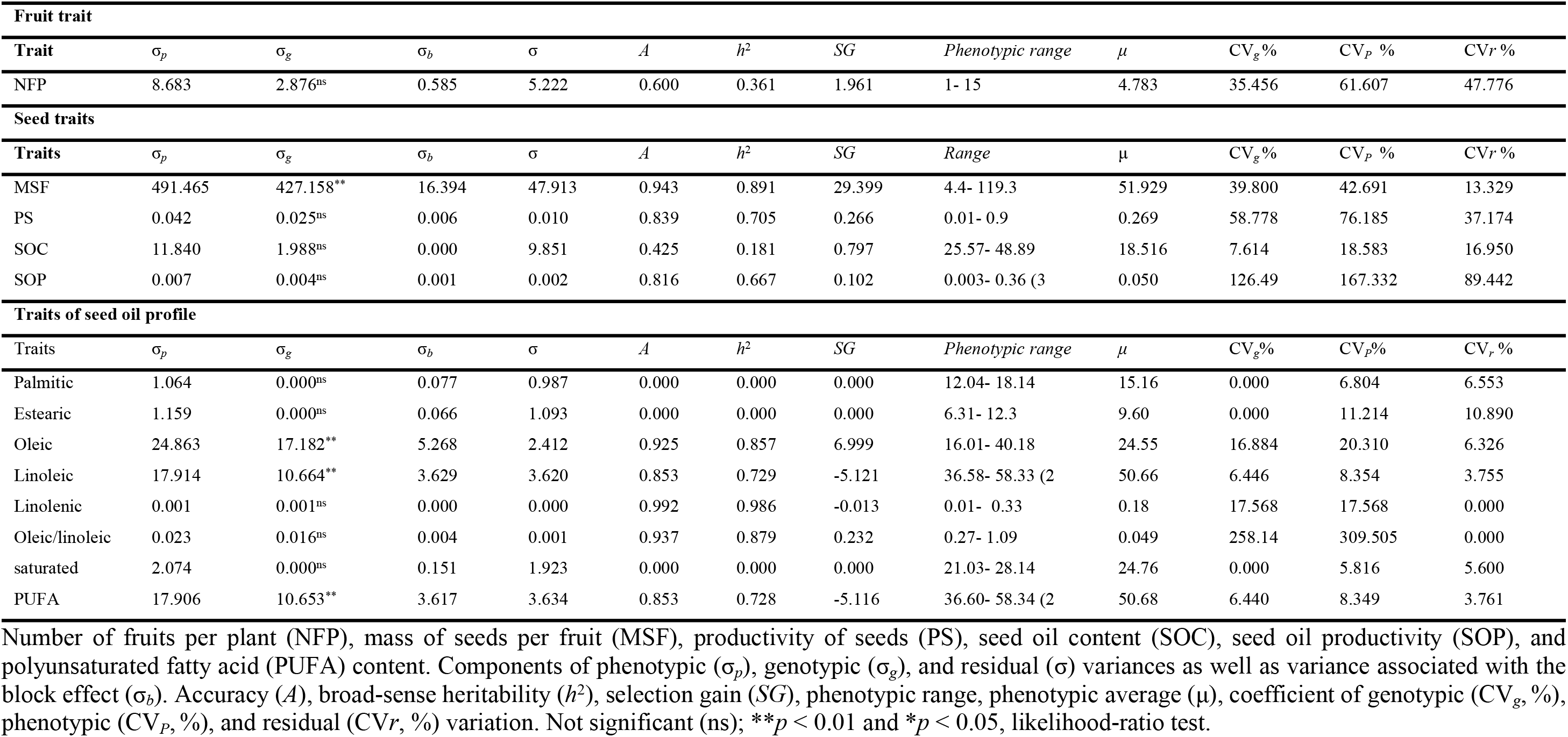
Variance components and genetic-statistical parameters of characteristics related to seed oil productivity and fatty acid profiles

Most of the tested characteristics showed very high heritability (>0.70) (Table 1), according to the classification of Resende (1995). The heritability estimates for PS, SOP, oleic acid content, and linoleic acid content were 0.705, 0.667, 0.857 and 0.729, respectively.

The predicted selection gains for PS and SOP were 0.266 and 0.10 t ha^-1^, respectively (Table 1). Among the characteristics related to oil fatty acid profile, oleic acid content achieved the greatest selection gain (6.99%), followed by linoleic acid content (−5.12%) and PUFA content (−5.11%), which also achieved considerable selection gains (Table 1). The phenotypic amplitude was up to 0.90 t ha^-1^ (phenotypic mean, 0.26 t ha^-1^) for PS and up to 0.36 t ha^-1^ (phenotypic mean, 0.050 t ha^-1^) for SOP (Table 1). Oleic acid content ranged from 16.01 to 40.18% (phenotypic mean, 24.55%), and linoleic acid content from 36.58 to 58.33% (phenotypic mean, 50.66%).

### 3.2 Correlations of characteristics related to SOP and fatty acid profiles

For convenience, NFP, MSF, and PS were assumed to be directly related to SOP. The phenotypic correlations of these first variables with SOP ranged from 0.51 (SOP × MSF) to 0.99 (SOP × PS), with all correlations being significant (*p* < 0.01). The phenotypic correlations between SOP × PS (0.99) and SOP × NFP (0.75) were classified as very strong and strong, respectively, according to Shimakura and Ribeiro Júnior (2012). The genotypic correlations between characteristics directly related to SOP were similar to the phenotypic correlations in terms of direction and significance, although their magnitude was slightly smaller (Figure 2).

**Figure 2.**
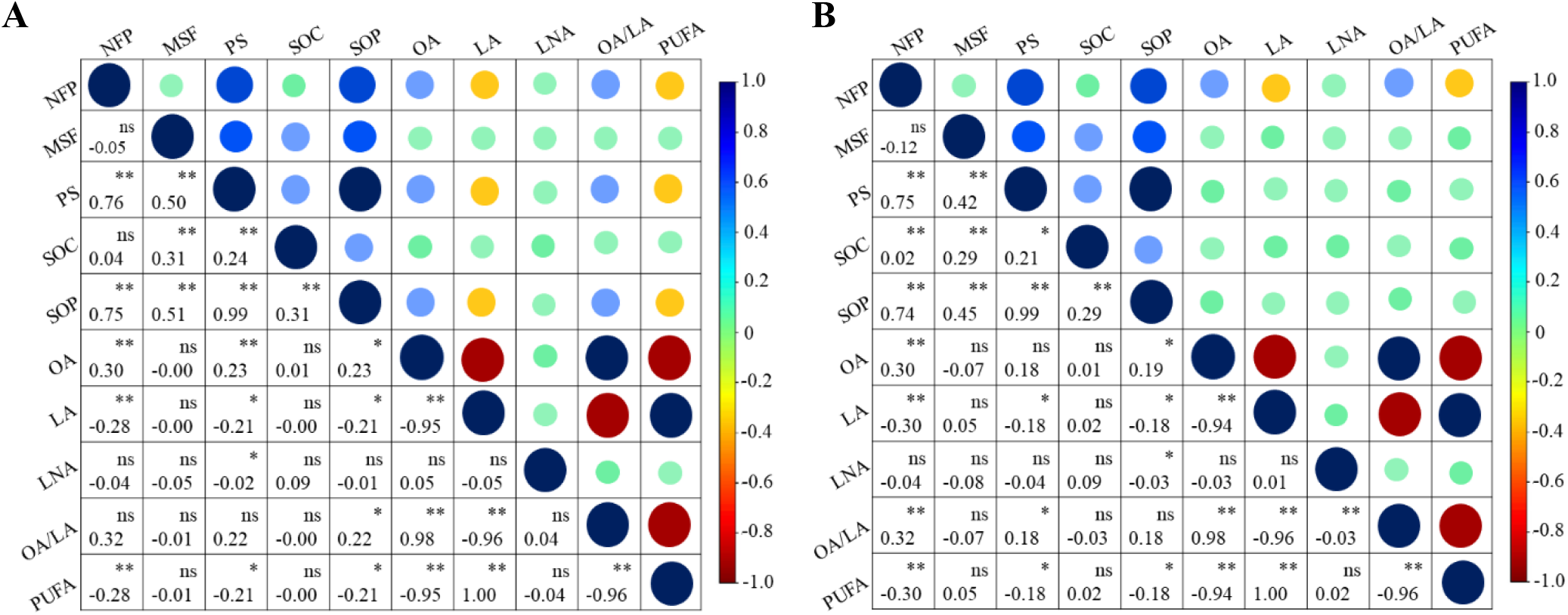
Phenotypic (A) and genotypic (B) correlations of characteristics related to seed oil productivity and fatty acids. Number of fruits per plant (NFP), mass of seeds per fruit (MSF), productivity of seeds (PS), seed oil content (SOC), seed oil productivity (SOP), oleic acid (OA) content, linoleic acid (LA) content, linolenic acid (LNA) content, OA:LA ratio, and polyunsaturated fatty acid (PUFA) content were measured.

SOP showed the strongest positive correlation with oleic acid content (0.23) and oleic acid:linoleic ratio (0.22) and strongest negative correlations with linoleic (−0.21) and PUFA (−0.21) content. SOP showed weak correlations with fatty acids, according to Shimakura and Ribeiro Júnior (2012), ranging from 0.20 to 0.39. Most of the genotypic correlations between SOP and oil fatty acids were similar to the phenotypic correlations in terms of direction and significance, although their magnitude was smaller (Figure 2).

Among fatty acids, linoleic and PUFA content showed the strongest positive correlation (1; *p* <0.01) (Figure 2). The correlation between the oleic content × oleic acid:linoleic ratio (0.98; *p* < 0.01) was classified as very strong (Figure 2), according to Shimakura and Ribeiro Júnior (2012). Oleic acid:linoleic ratio × linoleic acid content and oleic acid:linoleic ratio × PUFA acid content showed very strong negative correlations (both −0.96; *p* < 0.01). Similarly, oleic acid content × linoleic acid content showed very strong negative correlation (−0.95; *p* <0.01). Most of the genotypic correlations between fatty acids were similar to the phenotypic correlations in terms of direction, significance, and magnitude (Figure 2).

### 3.3 Clustering and variability of accessions based on characteristics related to SOP and fatty acid profiles

In clustering analysis, the tested accessions and controls formed five groups. Group 1 included 83 accessions (91.20%) and the controls Jabras, Jacarezinho, Maranhão, and Tetsukabuto. Group 2 comprised eight accessions (8.79%), corresponding to the second largest group. Group 3 comprised two accessions, and the remaining groups (4 and 5) comprised a single accession each (Table 2).

**Table 2.**
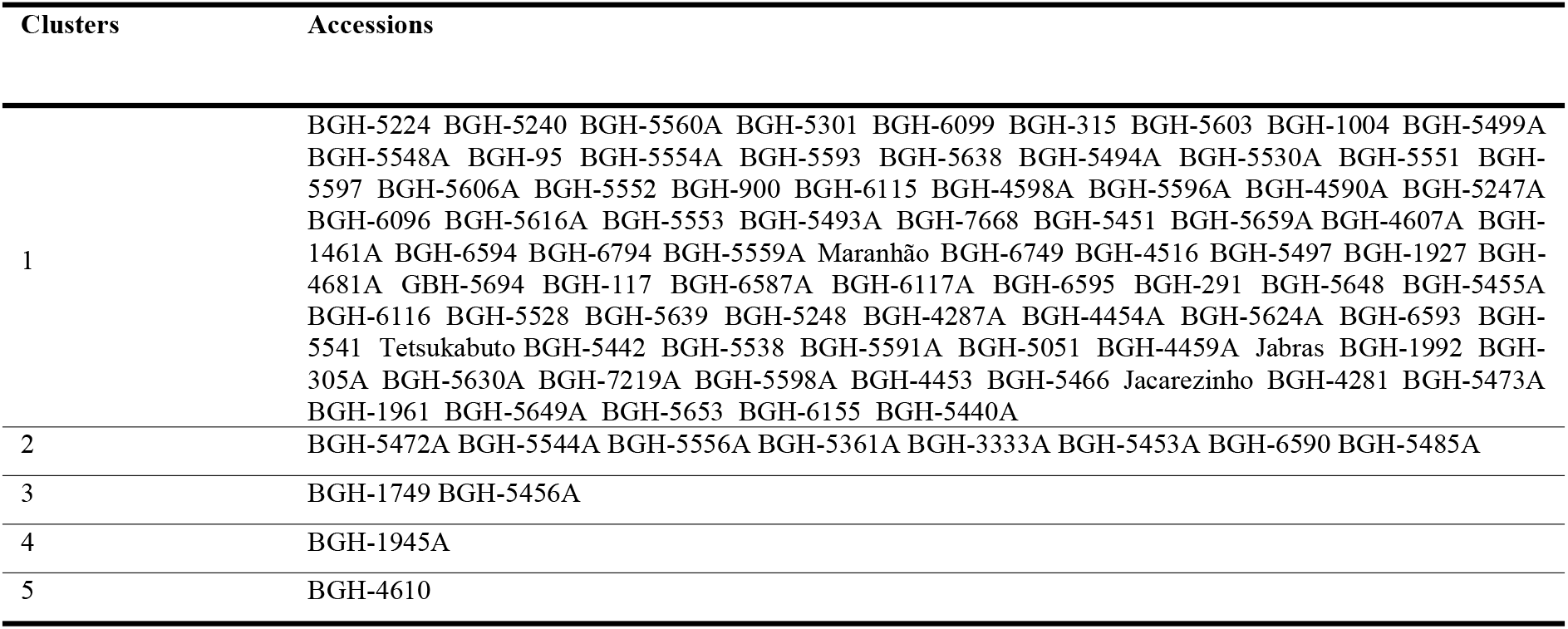
Tocher’s clustering of accessions based on the genotypic values of characteristics related to seed oil productivity and oil fatty acid profiles

| Clusters | Accessions |
| --- | --- |
| 1 | BGH-5224 BGH-5240 BGH-5560A BGH-5301 BGH-6099 BGH-315 BGH-5603 BGH-1004 BGH-5499A BGH-5548A BGH-95 BGH-5554A BGH-5593 BGH-5638 BGH-5494A BGH-5530A BGH-5551 BGH-5597 BGH-5606A BGH-5552 BGH-900 BGH-6115 BGH-4598A BGH-5596A BGH-4590A BGH-5247A BGH-6096 BGH-5616A BGH-5553 BGH-5493A BGH-7668 BGH-5451 BGH-5659A BGH-4607A BGH-1461A BGH-6594 BGH-6794 BGH-5559A Maranhão BGH-6749 BGH-4516 BGH-5497 BGH-1927 BGH-4681A BGH-5694 BGH-117 BGH-6587A BGH-6117A BGH-6595 BGH-291 BGH-5648 BGH-5455A BGH-6116 BGH-5528 BGH-5639 BGH-5248 BGH-4287A BGH-4454A BGH-5624A BGH-6593 BGH-5541 Tetsukabuto BGH-5442 BGH-5538 BGH-5591A BGH-5051 BGH-4459A Jabras BGH-1992 BGH-305A BGH-5630A BGH-7219A BGH-5598A BGH-4453 BGH-5466 Jacarezinho BGH-4281 BGH-5473A BGH-1961 BGH-5649A BGH-5653 BGH-6155 BGH-5440A |
| 2 | BGH-5472A BGH-5544A BGH-5556A BGH-5361A BGH-3333A BGH-5453A BGH-6590 BGH-5485A |
| 3 | BGH-1749 BGH-5456A |
| 4 | BGH-1945A |
| 5 | BGH-4610 |

Group 5, formed by the accession BGH-4610A, presented the highest average (μ+g) value for SOP, estimated at 0.277 t ha^-1^. Group 2, formed by the accessions BGH-5472A, BGH-5544A, BGH-5556A, BGH-5361A, BGH-5453A, BGH-3333A, BGH-6590, and BGH-5485A, also presented a high average (μ+g) value for SOP, estimated at 0.19 t ha^-1^ (Tables 2 and 3).

**Table 3.**
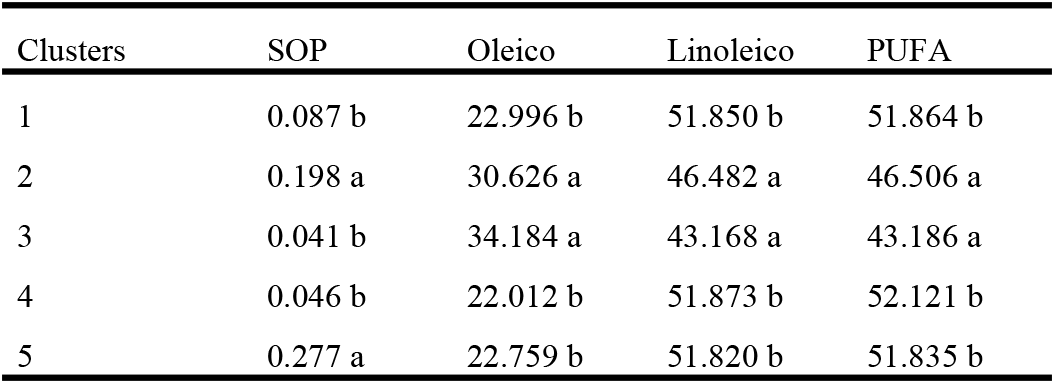
Grouping of genotypic averages of the accession groups based on Tocher’s method of average grouping

Regarding the fatty acid content of seed oil, group 3, formed by the accessions BGH-5456A and BGH-1749, presented the highest average (μ+g) value for oleic acid content (34.18%), followed by group 2 (30.62%). Group 3, formed by the accessions BGH-5456A and BGH-1749, presented the lowest average (μ+g) value for linoleic acid content (43.16%) (Tables 2 and 3).

The genotypic averages followed by the same letters in the column do not differ from one another based on the Tocher’s method of average grouping.

### 3.4 Principal components analysis

The first three principal components (PCs) explained 82.13% of total variation among accession in terms of characteristics related to SOP and fatty acid profiles. PC1 explained 47.16% of total variation. The oleic acid:linoleic acid ratio and oleic acid content were the characteristics with the highest positive loading, while linoleic acid content and PUFA content were the characteristics with the highest negative loading on PC1 (Figure 3).

**Figure 3.**
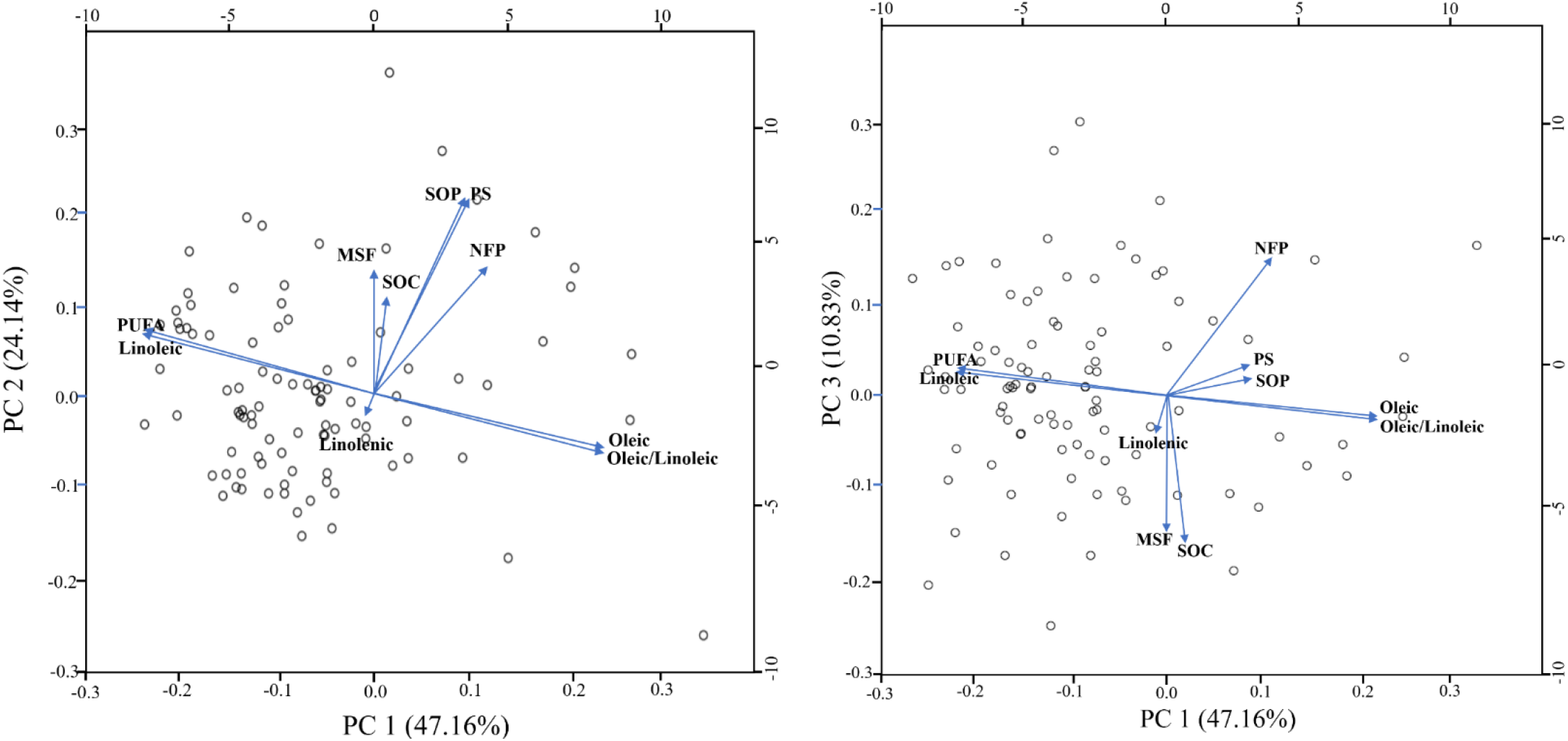
Dispersion of characteristics related to seed oil productivity and fatty acid profile in relation to the first three principal components. Number of fruits per plant (NFP), mass of seeds per fruit (MSF), productivity of seeds (PS), seed oil content (SOC), seed oil productivity (SOP), and polyunsaturated fatty acid (PUFA).

PC2 explained 24.14% of total variation, and PS, SOP, and MSF were the characteristics with the highest positive loading on this PC. PC3 explained 10.83% of total variation, and SOC and MSF were the characteristics with the highest loading on this PC.

Furthermore, PCA revealed the relationships between the characteristics. Along PC1, SOP was strongly and positively correlated with PS and NFP; moreover, along the same PC, there was a strong positive correlation between oleic acid content and oleic acid:linoleic acid ratio and a strong negative correlation between linoleic acid content and PUFA content (Figure 3).

### 3.5 Identification of promising accessions in terms of SOP and fatty acid profiles

Among the tested accessions, the (μ+g) estimate for SOP ranged from 0.14 to 0.27 t ha^-1^, which was much higher than the general average for accessions (0.05 t ha^-1^). Notably, the accessions BGH-4610A, BGH-5485A, BGH-6590, BGH-5556A, BGH-5472A, and BGH-5544A were the most promising in terms of SOP, with values close to 0.20 t ha^-1^ (Table 4).

**Table 4.**
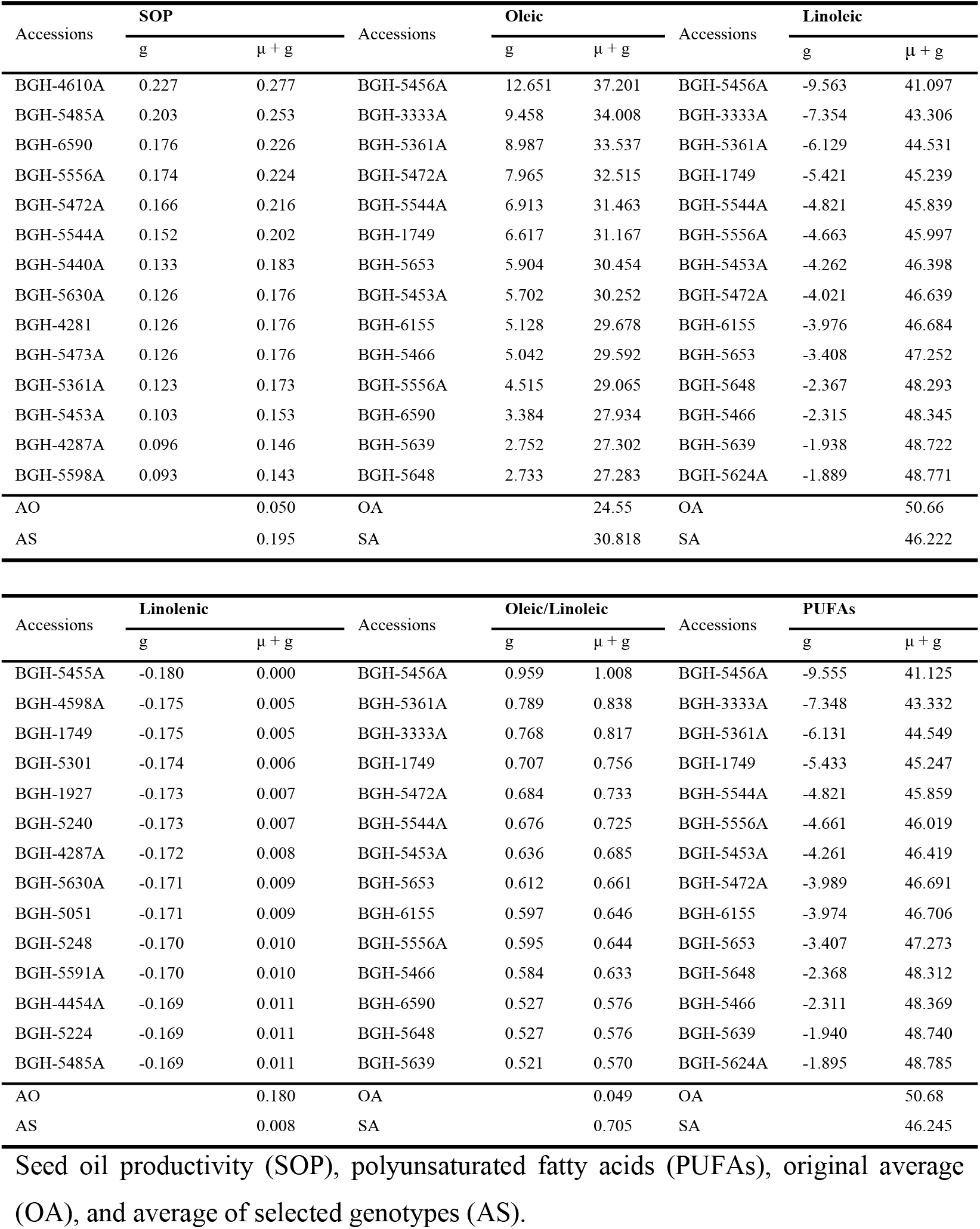
Estimates of genotypic values (g) and environmental interaction-free genotypic values (μ+g) for seed oil productivity and fatty acid profiles

Among fatty acids of seed oil, oleic acid content showed the highest (μ+g) amplitude (17.71 to 37.20%) (Figure 4). Associated with this, the (u+g) estimates for oleic acid content among the selected accessions ranged from 27.28 to 37.20% (Table 4). The accessions BGH-5456A, BGH-3333A, BGH-5361A, BGH-5472A, BGH-5544A, BGH-1749, BGH-5653, and BGH-5453A were the most promising in terms of oleic acid content, with values close to 30.00%.

**Figure 4.**
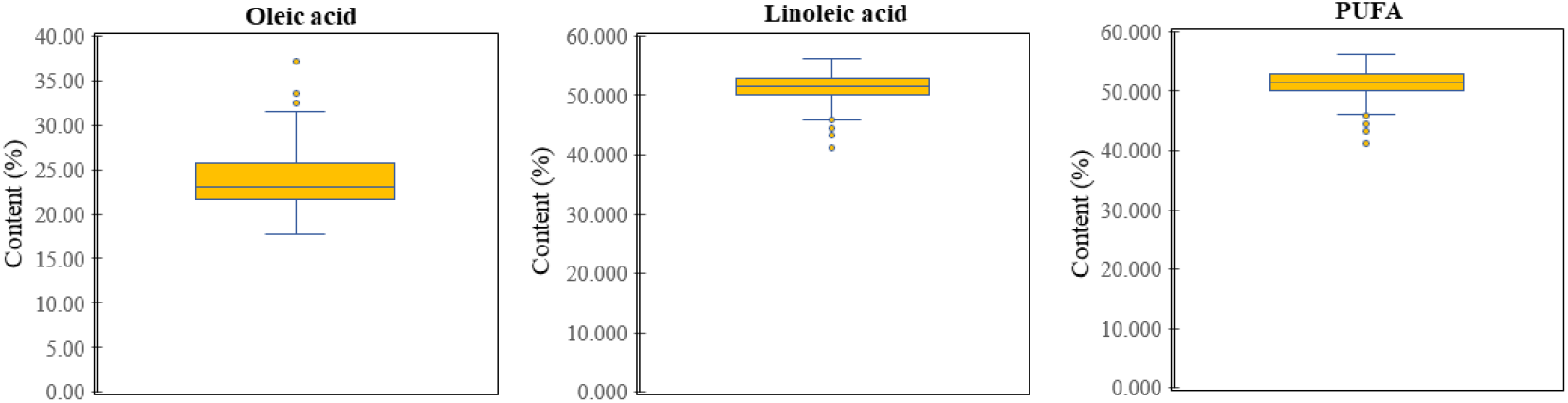
Box plots showing the variability in the oleic, linoleic, and polyunsaturated fatty acid content in the seed oil of the tested accessions.

Among the selected accessions, the (μ+g) estimate for linoleic content ranged from 41.09 to 48.77%. The accessions BGH-5456A, BGH-3333A, BGH-5361A, BGH-1749, BGH-5544A, and BGH-5556A expressed the lowest estimates for linoleic acid content, with values close to 40.00% (Table 4).

The (μ+g) estimates for oleic acid:linoleic acid ratio ranged from 1.00 to 0.57 among the selected accessions, and the accessions BGH-5456A, BGH-5361A, BGH-3333A, BGH-1749, BGH-5472A, and BGH-5544A expressed the lowest estimates, with values close to 0.75. The (μ+g) estimates for PUFA content ranged from 41.12 to 48.78%, and the accessions with the lowest estimates were BGH-5456A, BGH-3333A, BGH-5361A, BGH-1749, and BGH-5544A, with values close to 40% (Table 4).

## 4. DISCUSSION

### 4.1 Variance components and genetic-statistical parameters of characteristics related to SOP and fatty acid profiles

The greatest genotypic variances for MSF and PS, which were associated with their high heritability (0.89% and 0.70%, respectively), confirmed the marked genetic variability in these two characteristics, particularly for MSF, in the germplasm (Table 1). From these results, the predicted selection gains were significant and up to 29.39 g for MSF and 0.26 t·ha^-1^ for PS (Table 1). Consistent with our results, remarkable variability in characteristics related to seed production, such as MSF and PS, in the *C. moschata* germplasm has been reported previously (Lima Neto, 2013; Darrudi et al., 2018; Oliveira et al., 2020).

Regarding the fatty acid profile of seed oil, the highest genotypic variance and very high heritability for the content of oleic acid, demonstrate the high genetic variability and feasibility of identifying *C. moschata* accessions that can produce oil with higher oleic acid content. As a result, the predicted selection gain for oleic acid was 6.99%, which corresponded to the greatest selection gain among the components of seed oil (Table 1). Furthermore, the linoleic acid and PUFA content exhibited high genetic variability, also demonstrating the feasibility of identifying *C. moschata* accessions that can produce oil with lower linoleic acid content, providing predicted selection gains of −5.12% for this first fatty acid (Table 1).

The results for the fatty acid profiles obtained in the present study are consistent with previously reported profiles (Sobreira, 2013; Jarret et al., 2013). For instance, Jarret et al. (2013) evaluated the fatty acid profile of 38 seed samples corresponding to *C. moschata* accessions in the Plant Germplasm Collection of the United States Department of Agriculture, Griffin, and reported that oleic acid presented the greatest amplitude (10 to 53.80%), followed by linoleic acid (24.70 to 61.70%). Previous studies also corroborate high linoleic and oleic acid content, low palmitic acid content, and trace stearic and linolenic acid content in *C. moschata* seed oil (Applequist et al., 2006; Kim et al., 2012; Veronezi and Jorge, 2015). To the best of our knowledge, however, no previous study on *C. moschata* has analyzed the genetic parameters associated with the components of the fatty acid profile of seed oil, such as genetic variance, heritability, and selection gain.

Studies on other oilseeds, such as soybean, rapeseed, and sunflower, have often reported variations in the fatty acid profiles among germplasm samples and attributed such variations to genetic factors (Hemingway, et al., 2015; Yol et al., 2017). Given the quantitative nature of the fatty acid profile, it is reasonable to assume that this characteristic is also influenced by environmental factors, and some studies have shown the strong influence of environmental factors, such as temperature, on the fatty acid profiles of various oilseeds (Werteker et al., 2010). Corroborating the findings of Gomes et al. (2020), most of the accessions evaluated in the present study were acquired from family farmers, who do not typically select accessions for the characteristics of seed or seed oil, which possibly contributed to the maintenance of high variability in these traits.

### 4.2 Correlations of characteristics related to SOP and fatty acid profiles

The phenotypic correlations of NFP, MSF, and PS with SOP ranged from moderate to very strong (*p* < 0.001), corroborating the notion that these characteristics are directly related to SOP (Figure 2). The genotypic correlations of these first three variables with SOP were very similar to the phenotypic correlations, indicating that the relationship of the variables NFP, MSF, and PS with SOP may be attributed to genetic factors (Figure 2). These trends were repeated for correlations of other variables, which tended to express genotypic correlations similar to the phenotypic.

In the present study, we expected the variables NFP, MSF, and PS to be strongly correlated with SOP, consistent with previous reports of strong correlation between seed yield and oil productivity (Assefa et al., 2018). Thus, in addition to PS and MSF, NFP appears to be a determinant of greater SOP in *C. moschata*. These correlations of SOP with its directly associated characteristics can be strategic to guide indirect selection aimed at increasing *C. moschata* oil productivity. Thus, the selection of genotypes with higher PS and MSF may be a promising alternative to obtain greater SOP in *C. moschata*.

The positive correlations of SOP with oleic acid content and oleic acid:linoleic acid ratio observed in the present study suggest the feasibility of obtaining greater oil productivity while maintaining a desirable oil profile, with high oleic acid content and oleic acid:linoleic acid ratio (Figure 2). Additionally, the negative correlations of SOP with linoleic acid and PUFA content indicate the feasibility of obtaining greater SOP while maintaining lower PUFA such as the linoleic acid. To the best of our knowledge, no study on *C. moschata* has addressed the relationship between SOP and components of seed oil fatty acid profile. Meanwhile, studies on other oilseeds, such as peanut, have reported weak phenotypic correlations between SOP and oleic acid content (−0.034) (Yol et al., 2017), suggesting a small change in the acid content with increase in oil productivity in this crop. In contrast to the present study, Yol et al. (2017) reported a weak but positive correlation between SOP and oleic acid content.

Regarding the correlations among the components of oil fatty acid profile, the strong, negative but significant correlations of oleic acid content with linoleic acid and PUFA content indicate an antagonistic relationship between these components of *C. moschata* seed oil (Figure 2). The phenotypic correlations among these characteristics may be attributed to environmental and genetic factors. For instance, environmental factors, such as temperature, may strongly influence the correlations among fatty acids such as oleic and linoleic acid. In line with this, studies on other oilseeds, such as soybean, have shown that the oleic acid content of oil tends to decrease with increasing temperature, contrary to that observed for linoleic acid content (Bachlava et al., 2008), corroborating the negative correlations among these two fatty acids observed in the present study.

The metabolic pathways of fatty acids may be the key genetic factor responsible for the correlations among the components of oil fatty acid profile. In this context, desaturases in the plastids and endoplasmic reticulum play a central role in fatty acid synthesis and catalyze their conversion to MUFAs or PUFAs (Long et al., 2018). Among these enzymes, delta-12 fatty acid desaturase 2 (Δ12-FAD2) converts oleic acid precursors into linoleic acid precursors (Ohlrogge and Browse, 1995), and according to Dehghan and Yarizade (2014), the *FAD2* gene family is rather ubiquitous and diverse in plants. Thus, the strong negative correlation between oleic and linoleic acid content observed in the present study may also be related to the action of FAD2 during the biosynthesis of these two fatty acids.

The analysis of correlations among characteristics is an important subsidy for plant breeding, which must contemplate several variables simultaneously (Dias et al., 2017), proving very useful when determining selection strategies. As shown in the present study, the plant germplasms maintained in banks commonly constitute a representative sample of the species gene pool, thus providing comprehensive information on the relationships among germplasm characteristics, which makes the correlation analysis very useful during the initial evaluation of plant germplasms conserved in banks. The information on correlations among the components of seed oil observed in the present study may be particularly meaningful in breeding programs aimed at improving the fatty acid profile of *C. moschata* seed oil, considering the feasibility of increasing oleic acid content while decreasing PUFA content, given the strong negative correlation between these two components.

### 4.3 Clustering and variability of accessions based on characteristics related to SOP and fatty acid profiles

The clustering of accessions confirmed the variability in characteristics related to SOP and fatty acid profiles among the studied accessions (Table 2). This clustering is consistent with the high estimates of genotypic variance and heritability for the evaluated characteristics related to SOP, including MSF and PS, as well as those related to the fatty acid profile of seed oil, including oleic and linoleic acid content (Table 1). The variability observed among the accessions tested in the present study also corroborates previous reports emphasizing remarkable variability in both agromorphological and molecular characteristics in the *C. moschata* germplasm (Ferriol et al., 2004; Barboza et al., 2012; Ferreira et al., 2016).

The clustering of accessions in the present study did not reflect a greater similarity between accessions from the same state or geographic region, consistent with the reports of agromorphological characteristics of the *C. moschata* germplasm from different geographic regions (Moura, 2003; Gomes et al., 2020). Unlike the present study, previous studies involving the analysis of fatty acid profile of the seeds of other oilseed crops, such as soybean, have reported that germplasms from regions at higher latitudes tended to express higher palmitic, stearic, and oleic acid content than germplasms from lower latitudes (Wu et al., 2017; Abdelghany et al., 2020), suggesting a greater similarity between the germplasms from the same region. In this line, Bachlava et al. (2008) observed that the oleic acid content of soybean tended to increase from lower to higher latitudes, indicating increase in the content of this fatty acid with decrease in temperature, contrary to the trends for linoleic and linolenic acid. Similarly, Song et al. (2016) observed that the oleic acid content of soybean seeds was negatively correlated with the duration of sunlight incidence. Thus, the greater similarity between the soybean germplasms from the same region may reflect regional ecogeographic characteristics; different from the results of the present study.

Already present in the diet of native peoples (Piperno et al., 2003; Dillehay et al., 2007), *C. moschata* is widely cultivated in Latin America, which is an important center of diversity for this vegetable. In addition, previous studies have highlighted the variability in the Brazilian germplasm of *C. moschata* (De Lima et al., 2016; Ferreira et al., 2016; Gomes et al., 2020), possibly as a result of the adaptation of this germplasm to a wide ecological range, constituted by diverse edaphoclimatic conditions. Additionally, the intrinsic characteristics of *C. moschata*, such as the occurrence of natural hybridization across populations, associated with the processes of selection and seed exchange practiced by populations involved in its cultivation, also contribute to the variability in the Brazilian germplasm of this vegetable (Gomes et al., 2020).

The similarity between accessions of different geographic regions observed in the present study suggests the adaptability of the germplasm, indicating the feasibility of its cultivation under edaphoclimatic conditions different from those in its regions of origin. Compared with other crops, such as soybean (Abdelghany et al., 2020), the possible adaptability of *C. moschata* germplasm observed in the present study represents an opportunity to meet the diverse demands of various cultivation regions and systems of this vegetable, specifically in Brazil, which is characterized by its continental dimension.

Additionally, crossbreeding between divergent genotypes has been conveniently explored in *C. moschata*, aiming at the exploitation of hybrid vigor for aspects related to growth habit, fruit and seed production, as well as for fruit chemical–nutritional aspects (El-Tahawey et al., 2015; Kumar et al., 2018). Based on genotypic data and environmental interaction-free genotypic values, our results of variability analysis will be particularly useful to assist the crossing of promising genotypes aimed at the exploration of hybrid vigor for characteristics related to SOP and fatty acid profiles.

### 4.4 Principal components analysis

In consonance with their respective variance components and genetic parameters (Table 1), the oleic acid, linoleic acid, and PUFA content made the greatest contributions to the discrimination of the genotypes in PCA, confirming the high variability of these characteristics (Figure 3). The results of PCA regarding the variability of accessions were consistent with the results of clustering analysis (Table 2), also corroborating the strong positive correlations of PS with SOP, as well as the positive correlation of SOP with the oleic acid content (Figure 2).

### 4.5 Identification of promising accessions in terms of SOP and fatty acid profiles

Based on their highest genotypic averages for SOP, the groups 5 and 2 were identified as the most promising for this characteristic (Table 3). Consistent with this result, *per se* analysis identified accession BGH-4610 from group 5 as the most promising in terms of SOP, with the (μ+g) estimate of 0.27 t ha^-1^ (Table 4). In addition, accessions BGH-5485A, BGH-6590, BGH-5556A, BGH-5472A, and BGH-5544A from group 2 were also identified as promising in terms of SOP in *per se* analysis, with the (μ+g) estimate of ~0.20 t ha^-1^ (Table 4).

The groups 3 and 2 were identified as the most promising in terms of SOP with higher oleic acid content (Table 4). Associated with this, the accession BGH-5456A from group 3 was identified as most promising in terms of high oleic acid content in *per se* analysis, with the (μ+g) estimate of 37.20% (Table 4). The accessions BGH-3333A, BGH-5361A, BGH-5472A, BGH-5544A, and BGH-5453A from group 2 and the accession BGH-1749 from group 3 were also identified as promising in terms of high oleic acid content, with the (μ+g) estimate of ~30.00% (Table 4). The identification of promising groups and *per se* identification of accessions with high oleic acid content in the seed oil are associated with the high amplitude of (μ+g) estimates for this characteristic (Figure 4).

The groups 3 and 2 also presented the lowest averages for linoleic acid and PUFA content, confirming them as the most promising in terms SOP with lower PUFA content (Table 4). Consistent with this result, the accession BGH-5456A, BGH-3333A, BGH-5361A, BGH-5544A, BGH-5556A, and BGH-1749 were identified as promising in terms of low linoleic acid content, with the (μ+g) estimate of ~45% (Table 4). In *per se* analysis, the accessions identified as the most promising in terms of low linoleic acid also were the most promising in terms of low PUFA content, demonstrating the predominance of linoleic acid among PUFAs.

Recently, the development of cultivars with an oil profile that is better suited to human nutrition and health has been emphasized in the genetic breeding of oilseed crops. This is in agreement with a series of studies demonstrating the association between the consumption of lipid sources predominantly comprising saturated fatty acids and the high risk of cardiometabolic pathologies, particularly cardiovascular diseases and type II diabetes mellitus (Harris et al., 2009; Keys et al., 2017; Wu et al., 2019). This has encouraged the replacement of saturated lipids in human food by unsaturated fatty acids, with a particular focus on vegetable oils—the main source of unsaturated fatty acids in the human diet.

Associated with high levels of unsaturated fatty acids, vegetable oils should ideally have high stability against environmental stressors, such as humidity, light, heat, and oxygen. These oils must also be resistant to oxidative actions, which are related to the production of secondary components responsible for triggering allergic responses and cardiovascular diseases, such as atherosclerosis (Yanishlieva et al., 2001; Garbin et al., 2013), also responsible deteriorating the sensory quality (Choe and Min, 2006) and reduce the shelf life of oils (Xie et al., 2019). Given these demands, breeding programs for oilseed crops such as soybean, rapeseed, and corn have emphasized the development of cultivars that produce more stable oils, prioritizing the increase in the content of oleic acid, a MUFA, which has greater oxidative stability (Burton et al., 2006; Bachlava et al., 2008; Wang et al., 2009; Long et al., 2018). In this line, some studies have confirmed the higher oxidative stability of oleic acid [C18: 1 (Δ^9^)] compared to PUFAs such as linoleic acid [C18: 2 (Δ^9, 12^] and linolenic acid [C18: 3 (Δ^9, 12, 15^], indicating that oleic acid is 10 times more stable than linoleic acid and 20 times more stable than linolenic acid (Liu et al., 1992). Thus, genetic breeding for improving the fatty acid profile of *C. moschata* seed oil should simultaneously target an increase in oleic acid content and decrease in polyunsaturated fatty acids, particularly linoleic acid content, aiming at greater nutritional and physicochemical quality of oil.

The identification of *C. moschata* accessions with high oleic acid content in the seed oil indicates the feasibility of identifying accessions that produce seed oil with higher nutritional quality and stability. Of note, *C. moschata* seed oil contains high levels of bioactive components, such as vitamin E and carotenoids (Veronezi and Jorge, 2012), which are important antioxidants in the human diet and protect the oil against oxidative processes. Despite of their high nutritional value, a large portion of seeds produced during *C. moschata* cultivation is still discarded (Li et al., 2019), particularly in Brazil. As highlighted by Gomes et al. (2020), the use of *C. moschata* seeds for oil production represents an alternative strategy to complement the diet, in addition to increasing the income of farmers involved in the production of this vegetable.

To the best of our knowledge, the present study is the first to analyze the SOP and fatty acid profile of *C. moschata* based on a relatively large number of accessions representative of different geographic regions of Brazil. Our results demonstrate the marked potential of some *C. moschata* accessions for oil production, as confirmed by the *per se* identification of accessions with high SOP and high oleic acid content in the oil. Our results corroborate the productive potential as well as the nutritional and physicochemical quality of seed oil from the Brazilian germplasm of *C. moschata*.

## 5. CONCLUSIOSNS

The tested accessions expressed high genetic variability in terms of MSF and PS, providing the predicted selection gains of 29.39 g and 0.26 ha^-1^, respectively.

Phenotypic and genotypic correlations indicated that a greater *C. moschata* SOP can be achieved by selecting for higher PS and MSF. Correlations also indicated that a greater SOP can be obtained while maintaining high oleic fatty acid content and low linoleic acid content, providing oil with better nutritional and chemical quality.

In the analysis of variability, the 91 accessions tested in this study were clustered into five groups, allowing the identification of the most promising groups in terms of greater SOP and higher oleic acid content in the oil, an approach that will guide the use of this germplasm in breeding programs aimed at improving the SOP and fatty acid profile.

*Per se* analysis identified the accessions BGH-4610, BGH-5485A, BGH-6590, BGH-5556A, BGH-5472A, and BGH-5544A as the most promising in terms of SOP, with the (μ+g) estimate of ~0.20 t ha^-1^. Accessions BGH-5456A, BGH-3333A, BGH-5361A, BGH-5472A, BGH-5544A, BGH-5453A, and BGH-1749 were identified as the most promising in terms of higher oleic content in oil were, with the (μ+g) estimate of ~30%, and most of these accessions were also the most promising in terms of lower linoleic acid content in oil, with the (μ+g) estimate ~40%. Therefore, part of the *C. moschata* germplasm evaluated in the present study is a promising source for the genetic improvement of SOP and fatty acid profile, aiming at the production of oil with better nutritional and physicochemical quality.

NFP: number of fruits per plant
MSF: total mass of seeds per fruit
PS: productivity of seeds
SOC: total seed oil content
SOP: seed oil productivity
UFV: Federal University of Viçosa
BLUPs: best non-biased linear predictions
BLUEs: best non-biased linear estimates
PUFAs: polyunsaturated fatty acids
RML: restricted maximum likelihood
MUFA: monounsaturated fatty acids

## AUTHOR CONTRIBUTIONS

### Conceptualization

Derly José Henriques da Silva.

### Data curation

Ronaldo Silva Gomes, Ronaldo Machado Júnior, Cleverson Freitas de Almeida, Rebeca Lourenço de Oliveira.

### Investigation

Ronaldo Silva Gomes, Ronaldo Machado Júnior, Cleverson Freitas de Almeida, Rebeca Lourenço de Oliveira, Ednângelo Duarte Pereira, Fabio Teixeira Delazari.

### Software

Rafael Ravaneli Chagas and Ronaldo Silva Gomes.

### Supervision

Derly José Henriques da Silva.

### Writing – original draft

Ronaldo Silva Gomes.

### Writing – review & editing

Ronaldo Silva Gomes, Derly José Henriques da Silva, Rebeca Lourenço de Oliveira, Cleverson Freitas de Almeida.

